# The differentiation of pluripotent stem cells towards transplantable endothelial progenitor cells

**DOI:** 10.1101/2021.02.05.430011

**Authors:** Kezhou Qin, Jun Yang

**Affiliations:** Department of Cell Biology, Institute of Basic Medical Sciences, Chinese Academy of Medical Sciences & Peking Union Medical College, 5 Dongdan San Tiao, Beijing 100005, China; Department of Physiology, and Department of Cardiology of the Second Affiliated Hospital, Zhejiang University School of Medicine, Hangzhou 310058, Zhejiang, China

**Keywords:** stem cell differentiation, endothelial cell, endothelial progenitor cell, pulmonary arterial hypertension

## Abstract

Endothelial progenitor cells (EPCs) and endothelial cells (ECs) have been applied in the clinic to treat pulmonary arterial hypertension (PAH), a disease characterized by disordered pulmonary vasculature. However, the lack of sufficient transplantable cells before the deterioration of disease condition is a current limitation to apply cell therapy in patients. It is necessary to differentiate pluripotent stem cells (PSCs) into EPCs and identify their characteristics. Comparing previously reported methods of human PSCs-derived ECs, we optimized a highly efficient differentiation protocol to obtain cells that match the phenotype of isolated EPCs from healthy donors. The protocol is compatible with chemically defined medium (CDM), it could produce a large number of clinically applicable cells with low cost. Moreover, we also found PSCs-derived EPCs express CD133, have some characteristics of mesenchymal stem cells and are capable of homing to repair blood vessels in zebrafish xenograft assays. In addition, we further revealed that IPAH PSCs-derived EPCs have higher expression of proliferation-related genes and lower expression of immune-related genes than normal EPCs and PSCs-derived EPCs through microarray analysis. In conclusion, we optimized a highly efficient differentiation protocol to obtain PSCs-derived EPCs with the phenotypic and molecular characteristics of EPCs from healthy donors which distinguished them from EPCs from PAH.

## 1. Introduction

In hypoxic environments, endothelial cells (ECs) lining on the inner layer of blood vessel have the ability to modulate vascular tone and are involved in angiogenesis (Niskanen et al. 2018). Endothelial progenitor cells (EPCs) are one type of ECs which could be isolated from blood by colony expansion and have been demonstrated to promote ischaemic tissue angiogenesis (Zhao et al. 2019), so they are capable of facilitating vascular repair under different ischaemic conditions, such as acute myocardial infarction, unstable angina, stroke, diabetic micro vasculopathies, pulmonary arterial hypertension, atherosclerosis, and ischaemic retinopathies (Ward, Stewart, and Kutryk 2007; Jung and Roh 2008; Sekiguchi, I1, and Losordo 2009). Human ECs can only be obtained from dissected vessel after surgical moving it from body. Although EPCs can be isolated by our group with improved technique from only 20 mL blood, they still have limited expansion potential as ECs, are rare in peripheral or umbilical cord blood compared with other blood cells, which hinders the application of cell therapy in clinical treatment (Medina et al. 2012; Medina et al. 2010; Zhang, Chu, et al. 2017).

Human PSCs including embryo stem cells (ESCs) and induced pluripotent stem cells (iPSCs) can be induced to produce scalable endothelial cells and endothelial progenitor cells for vascular remodelling (Belair et al. 2015; James et al. 2010; Lian et al. 2014; Medina et al. 2010; Nguyen et al. 2016; Patsch et al. 2015; Prasain et al. 2014), and so far there have been two methods of inducing PSCs to differentiate into vascular cells, namely, embryoid body formation (Park et al. 2014; Park et al. 2019) and monolayer-directed differentiation (Prasain et al. 2014; Patsch et al. 2015; James et al. 2010). In the former method, the cells need to be transferred into ultra-low-attachment plates to obtain embryoid bodies (EBs) to generate various types of cells, which is not cost-effective and is often time consuming (Han et al. 2018; Park et al. 2014). Monolayer differentiation methods have a higher differentiation efficiency (Patsch et al. 2015) than EBs methods, and have been the main methods for obtaining a large amount of functional ECs to repair vascular deficiencies, but it’ s necessary to further understand the complicated factors affecting EC differentiation.

Endothelial dysfunction has been thought to be the main contributor of PAH/IPAH. Some studies have reported that the numbers of CD133^+^ cells in peripheral blood increased in PAH/IPAH patients compared with controls (Foris et al. 2016; Toshner et al. 2009). Toshner et al. also demonstrated that PAH patient-derived endothelial progenitor cells have a hyperproliferative phenotype and a reduced capacity to form vascular networks (Toshner et al. 2009). However, recent results demonstrated that early EPCs overexpressing eNOS could be beneficial for the treatment of PAH (Gomberg-Maitland et al. 2013).

In this study, we aimed to explore the factors affecting vascular cell differentiation and to optimize a chemical method and cost-effective system for generating human PSCs-derived EPCs through monolayer-directed differentiation, while also characterizing the molecular features of PSCs-derived EPCs, IPAH-derived EPCs and normal EPCs. We found that the Rock inhibitor Y27632 and DMSO markedly accelerated vascular mesoderm generation from PSCs and improved the differentiation efficiency. Then, the functions of these cells were tested in parallel by tube formation and LDL uptake assays, microarray analysis and cell transplantation in zebrafish. This optimized system can offer a simple, rational, cost-effective platform to produce PSCs-derived ECs/EPCs for vascular research and clinical application.

## 2. Methods

### 2.1 Cell Maintenance

Human ESCs (H9 and H1, WiCell Madison, WI, given by Roger Pedersen, University of Cambridge) /iPSCs(generated in house, reported previously by Fang, Zhou) were cultured in Essential 8 (E8) medium or mTeSR™1 Complete Kit (Catalog #85850) or hPSC-CDM™(Cauliscell Inc. #400105) supplemented with hPSC-CDM™ supplement (Cauliscell Inc. #600301) on Matrigel-coated (BD Bioscences, #356230) 6-well plates and were passaged with 500 μM EDTA for 3-5min. EPCs/ECs were maintained in EGM2+16%FBS (HyClone) (Ormiston et al. 2015).

### 2.2 Endothelial Cell Differentiation, Purification, and Culture

When human ESCs/iPSCs grew to 80%-90% confluency, they were dissociated with Accutase (Gibco, #A11105-01) and plated about 3 x 10^4^ cells/well in vitronectin-coated (Cauliscell Inc.#500125) 12-well plates, ESCs/iPSCs were differentiated into mesoderm cells by culturingin E8 medium (Gibco, A1516901) supplemented with 25ng/mL Activin A(R&D, Catalog Number.338-AC), 10 μM Y27632 (Sigma) and 10ng/mL BMP4(R&D, Catalog Number.314-BP) for 3 days. Mesoderm cells were then cultured for 4 days in E6 medium (Gibco, A1516401) supplemented with FGF2 100ng/ml (R&D, Catalog Number.233-FB), VEGF 50ng/mL (R&D, Catalog Number.293-VE), BMP4 50ng/mL and SB431542 5μM(Sigma-Aldrich, CAS 301836-41-9-Calbiochem) to generate endothelial cells, and the whole differentiation process continues 7 days. Cells were counted and the cell suspension was prepared for the isolation of endothelial cells. Endothelial cells were isolated by using CD31^+^ MicroBeads (Miltenyi Biotec, Order no.130-091-935) according to manufacturer’ s instructions and cultured in EGM2 with 16% FBS (HyClone).

### 2.3 Flow cytometry

At day 3 or 7 of differentiation, adherent cells were harvested using 0.25% TrypleE with EDTA and made into a single-cell suspension in PBS with 0.2% BSA; then all cells were analyzed using flow cytometry directly without purification. Mouse Anti-Human APJ APC-conjugated Antibody(R&D, Catalog Number: FAB8561A) was used as a ratio of 1: 50, Mouse IgG1 (FITC, Material Number:551954, BD Pharmingen), Mouse Anti-human CD31 (CD31-FITC, Material Number: 555824, BD Pharmingen), Mouse IgG1-APC (Order no:130092214, Miltenyi Biotec), Mouse Anti-human CD34 (CD34-APC, Material Number: 560940, BD Pharmingen), Mouse Anti-human CD43 (CD43-APC, Material Number: 560198, BD Pharmingen), Mouse Anti-Hamster IgG PE-conjugated Antibody (Catalog Number: F0120,R&D system), Mouse Anti-human KDR (KDR-PE, FAB357P, R&D) and Mouse Anti-Human NRP-1 (NRP-1-PE, Material Number: 565951, BD) antibodies were used at a ratio of 1:20. Single-cell suspensions were subsequently incubated with antibody or antibodies at 4°C for about 40 mins. Flow cytometric detection of the cell surface antigens were performed on a BD Accuri™ C6 Plus personal flow cytometer (Becton Dickinson). Compensation was set by single-positive controls.

### 2.4 uptake of acetylated LDL

PSCs-derived ECs/EPCs were incubated with 10g/mL DiI-Ac-LDL (Molecular Probes) in serum-free EBM-2 (Lonza) for 4 hours at 37°C, respectively.

### 2.5 In vitro capillary network formation assay on Matrigel

PSCs-derived ECs/EPCs were trypsinized and resuspended in EGM-2 media with 16% FBS. Cells were plated at a density of 1.0 × 10^4^ cells per well in triplicate in 96-well plates coated with 50 μl of growth factor - reduced Matrigel (BD Biosciences). Plates were incubated overnight at 37° C. After 6h of incubation, photomicrographs were taken from each well at 10x magnification using an Olympus CKX41 microscope with a 10x objective.

### 2.6 Cell transplantation and therapy in Zebrafish

Zebrafish, the transgenic line Tg (Flk: GFP) were maintained according to standard procedures in compliance with local approval. After 48 hours post fertilization (hpf), embryos were used to inject PSCs-derived ECs/EPCs stained with CM-Dil at approximately 60μm above the ventral end of the duct of Cuvier, then the embryos were maintained at 30°C for 48 hours. Fluorescent image acquisition was performed using a Leica MZ16FA stereo-microscope. Zebrafish embryos were treated with sugen5416 for 24 hours at 4 hpf, then 20-30 fish were respectively therapied by PSCs-derived ECs/EPCs /medium (EGM2 + 16%FBS) and maintained at 30°C for 48 hours, the number of normal zebrafish were recorded. The experiments were repeated three times. Euthanasia of all zebrafish was performed by exposure to bleach or rapid freezing followed by maceration.

### 2.7 EPCs from adult peripheral blood samples

EPCs were isolated from human peripheral blood (PB) which were obtained from six female donors, including three IPAH patients and three normal adults, aged between 25 and 40 years. Fresh human PB (20mL) was obtained under full ethical approval; then mononuclear cells (MNCs) were isolated from PB by density gradient centrifugation and cultured in EGM2(Lonza) supplemented with 16% FBS.

### 2.8 RNA extraction, cDNA synthesis and RT-qPCR

Total RNA from human cell lines was extracted with Trizol (Life Technologies). RNA yield was determined using the NanoDrop ND-1000 spectrophotometer (NanoDrop Technologies). Total RNA (1 μg) was converted to cDNA using the PrimeScript™ RT reagent Kit with gDNA Eraser (TAKARA). Quantitative PCR (qPCR) was done using TransStart Tip Green qPCR SuperMix (TransGen Biotech) and detection was achieved using the CFX Connect™ Real-Time System (BIO-RAD). Primer sequence are listed in Supplementary Table 1. Expression of target genes was normalized to reference gene *GAPDH*.

### 2.9 Microarray analysis

Samples including IPAH-EPC1, IPAH-EPC2, IPAH-EPC3, normal EPCs (Con1, Con2, Con3) and PSCs-derived EPCs (H7EC, H9EC and 202EC) were used to do microarray analysis with HG-U133 Plus_2. Microarray data were carried out quality control, then normalized using rma method from R package affy. Differentially expressed genes (DEGs) were obtained by T test. DEGs (log FoldChange <-2 or >2, and p-Value <0.05) were carried out gene ontology (GO) clustering analysis by R package clusterProfiler (Yu et al. 2012).

### 2.10 Immunofluorescence assay

PSCs-derived ECs/EPCs were fixed with 4% (w/v) paraformaldehyde for 10 min and permeabilized with 0.1% (v/v) Triton X-100 in PBS for 5 min. After blocking with 10% (v/v) donkey serum (Solarbio, SL050) for 30 min, cells were incubated with primary following antibodies: anti-CD133 (ABclonal, 1155750301, 1:100) overnight at 4° C. Cells were washed with PBS, then incubated with secondary antibodies conjugated with Alexa-488 (Molecular Probe) and visualized by confocal microscopy. The confocal images were obtained with a Zeiss confocal microscope. All the images were taken at room temperature and images were analyzed using a ZEN 2.6(blue edition).

### 2.11 Statistical analysis

Statistical analyses were performed using Student’ s t-test and data were reported as mean ± s.d. or standard error of the mean, Figure legends show the number of biological repeats for each experiment (n), the experiments were repeated three or more than three times. Statistical significance was assumed when P <0.05.

## 3. Results

### 3.1 Exploring the factors affecting PSCs-derived EC differentiation efficiency

A few protocols for PSCs-derived ECs have been reported by others can generate about 20% ECs (Zhang, Schwartz, et al. 2017). However, it is still not enough for cell therapy or other transplantation purpose. So we tried to identify the factors affecting endothelial cell differentiation and optimize the protocol according to recent published protocols (Duan et al. 2018; Patsch et al. 2015; Wang et al. 2018) to provide a schematic diagram (Fig. 1A). Firstly, we seeded approximately 3 x 10^4^ cells/well in a 12-well plate instead of approximately 1 x 10^5^ cells/cm^2^ in previous studies. Secondly, we performed the differentiation with increasing cell number and found that the efficiency decreased with seeding cells number more than 10000 per well for H1 ESCs (Fig. 1B). Then, considering the high occurrence of PAH in females, we used H9-ESCs(female) for subsequent experiments. As an important mesoderm inducer, BMP4 was applied at an optimized concentration in our two-stage protocol (first stage and second stage). We also tested whether BMP4 affected PSCs-derived EC differentiation efficiency in the second stage and found that in the presence of BMP4 the CD31^+^ cells could be obtained about double the number than the combination without BMP4(Fig. 1C); BMP4 also promoted the generation of CD31^+^CD34^+^ cells at the same level (Bai et al. 2010). With the increased concentration of SB431542 (5 μM, 10 μM and 20 μM) in the second stage, the differentiation efficiency increased (Fig. 1D-E). Thirdly, mesoderm cells treated with Rock inhibitor were induced to differentiate into skeletal myocytes (Sadahiro et al. 2018), so we predicted that Y27632 plays the same role in the process of EC differentiation. In addition, we used Y27632 to improve cell survival after passage by treating 202-iPSCs with Y27632 for one day or three days and showed that the three-day induction resulted in a higher differentiation rate (41.9% ± 4.78%) than the one-day induction (6.37% ± 1.07%) (Fig. 1F-G). From the results above, we concluded that the differentiation efficiency decreases with increasing seeding cell number (within 10,000-60,000), BMP4 and SB431542 promote PSCs-derived EC differentiation in the second stage, and Y27632 promotes hESC and iPSC differentiation towards ECs. However, not all lines have the same increase rate as these hPSCs, especially in patient derived iPSCs, in the next step we further examine the EC differentiation potential with some reagents.

**Fig. 1.**
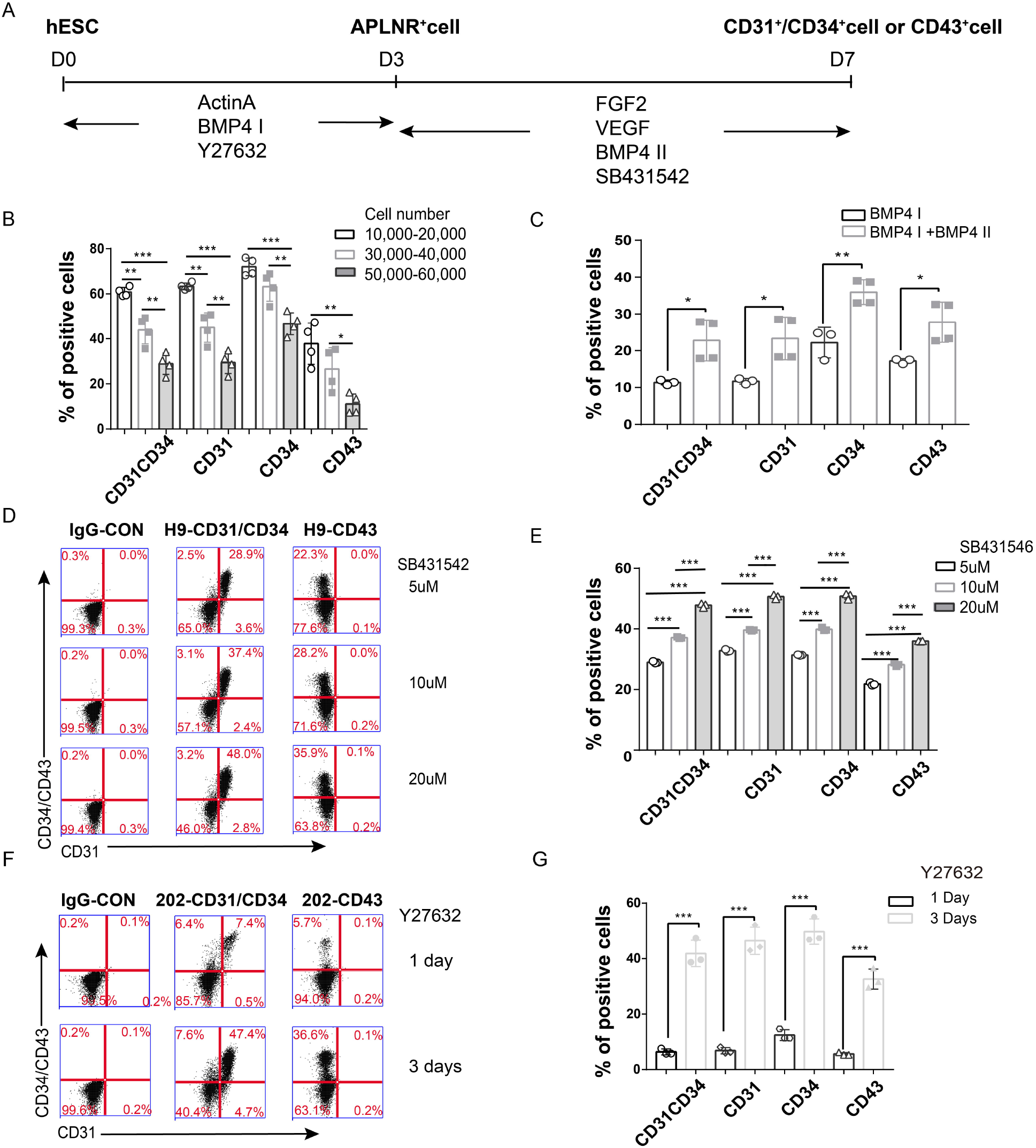
Identifying factors for PSCs-derived EPCs. (A) A schematic diagram of inducing human PSCs-derived ECs. (B) Number of seeding cells affects PSCs-derived EC differentiation efficiency as analysed by flow cytometry. (C) In the second step of PSCs-derived EC differentiation, adding BMP4 promotes PSCs-derived EC differentiation efficiency as analysed by flow cytometry. (D) and (E) In the second step of EC differentiation, appropriately increasing SB431542 promotes PSCs-derived EC differentiation efficiency as analysed by flow cytometry. (F) and (G) The inhibitor Y27632 improves PSCs-derived EC differentiation efficiency as analysed by flow cytometry in the first step of 202-iPSC-EC differentiation when Y27632 was used for three days. Statistics of CD31^+^CD34^+^ cells, CD31^+^ cells, CD34^+^ cells and CD43^+^ cells. Data are represented as the mean ± SD. n=3-4, the experiments were repeated more than three independent times. Student’s t test was performed (*p < 0.05, **p < 0.01, ***p < 0.001).

### 3.2 DMSO plays an important role in PSCs-derived hemogenic endothelial (HE) cell and PSCs-derived vascular EC differentiation

The differentiation of two main ECs type - hemogenic endothelial (HE) cell and vascular endothelial (VC) cell from human PSCs were both examined. We carefully compared the solvents of the reagents used in the differentiation process and found that when using Y27632 dissolved in DMSO, the differentiation efficiency was high, but when Y27632 was dissolved in ddH2O, the differentiation efficiency was low. Therefore, we hypothesized that the addition of DMSO results in increased differentiation efficiency. Also we observed higher differentiation rate when we supplied medium with Y27632 dissolved in DMSO rather than in H2O for one/three days (Fig. 2A-B). Furthermore, we obtained the same result in another chemically defined medium (CDM) (Supplementary Fig. 1). Therefore, DMSO is also an important reagent for PSCs-derived EC differentiation.

**Fig. 2.**
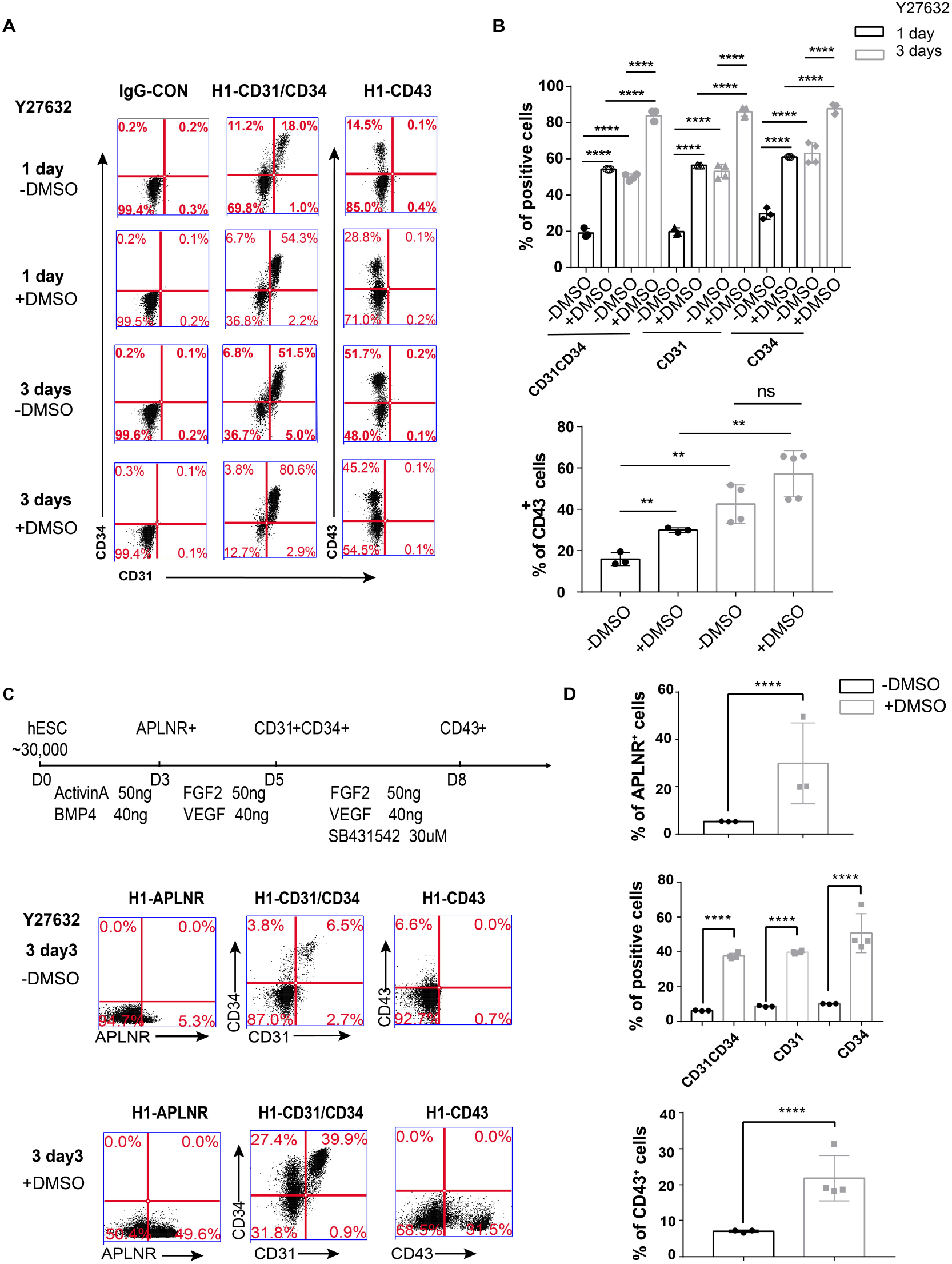
Adding DMSO improves EC and HE cell differentiation efficiency. (A) and (B) PSCs-derived EC (H1EC) differentiation efficiency increases when the cells are treated with DMSO in the first step for one day or three days. Statistics of CD31^+^CD34^+^ cells, CD31^+^ cells, CD34^+^ cells and CD43^+^ cells as analysed by flow cytometry. (C) and (D) Improved protocol for the HE differentiation of PSCs. A schematic digram of the stepwise induction process is shown, and DMSO treatment for three days in the first step promotes the differentiation efficiency as analysed by flow cytometry. Statistics of APLNR^+^ cells, CD31^+^CD34^+^ cells, CD31^+^ cells, CD34^+^ cells and CD43^+^ cells. Data are represented as the mean ± SD, n=3-4, and the experiments were repeated more than three times. Student’s t test was performed (ns: not significant, **p < 0.01, ****p < 0.0001).

We also improved the hemogenic endothelial cell differentiation protocol by adding Y27632 and DMSO in a previous protocol (Wang et al. 2018). When we added Y27632 in water and DMSO (1 μl/mL) for the first three days, we found that there was a higher differentiation efficiency than adding only Y27632 at all stages for APLNR^+^ cells (29.87%±17.09% versus 5.3%±0.1%), CD31^+^CD34^+^ cells (37.70% ± 1.55% versus 2.39%±0.17%), and CD43^+^ cells (21.85%±6.31% versus 9.17%±0.65%) (Fig. 2C-D). We also confirmed that adding DMSO promotes HE cell differentiation.

### 3.3 A modified protocol for generating ECs from human PSCs

Thus far, we have tested the factors affecting EC differentiation, including seeding cell number, induction time, BMP4, SB431542, Y27632 and DMSO to achieve best differentiation potential with H1 ESCs.

Then we seeded approximately 3 x 10^4^ cells/well in a 12-well plate using 10 μM Y27632 dissolved in DMSO and 10 ng/mL BMP4 for 3 days. After 7 days of differentiation, the cells were analysed directly using flow cytometry, which showed that H1 ESCs generated 92.17% ± 0.42% endothelial cells (Fig. 3A-D). In the end, we were able to optimize a highly efficient differentiation system of PSCs-derived ECs.

**Fig. 3.**
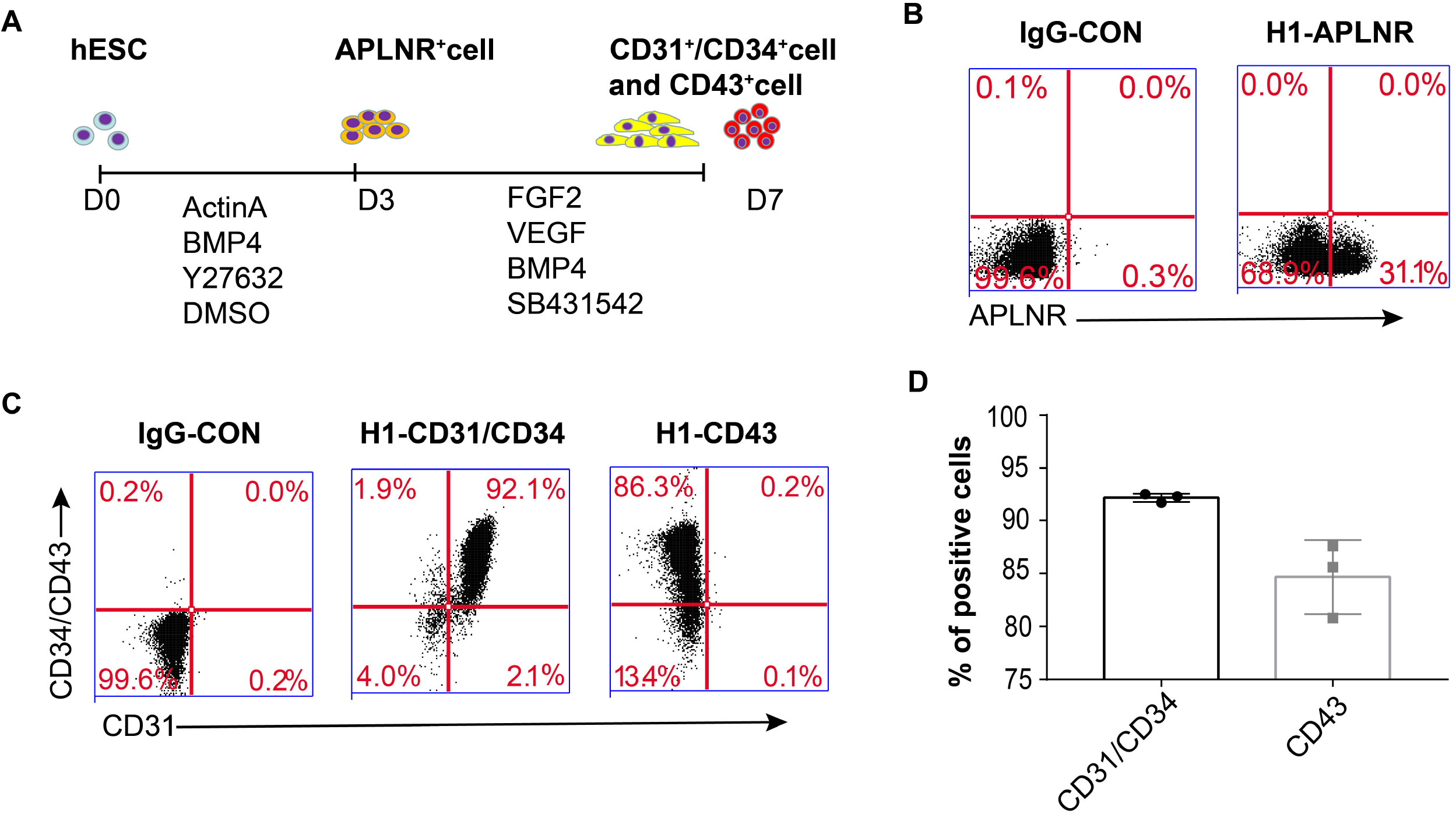
Improved protocol for the highly efficient differentiation rate of PSCs-derived ECs. (A) schematic diagram of inducing human PSCs-derived ECs via a mesoderm intermediate. (B) Representative yields of APLNR+ cells as analysed by flow cytometry. (C) Representative yields of CD34^+^CD31^+^ cells and CD43^+^ cells as analysed by flow cytometry after 7 days of differentiation. (D) Statistics of CD31^+^CD34^+^ and CD43^+^ cells. Data are represented as the mean ± SD, n=3, and the experiments were repeated three independent times.

### 3.4 PSCs-derived ECs function as endothelial cells

PSCs-derived ECs were isolated by using CD31^+^ MicroBeads and were cultured in EGM2 with 16% FBS. To characterize the PSCs-derived ECs, we carried out two assays, a Dil-Ac-LDL uptake assay and a tube formation assay. As a result, we verified that PSCs-derived ECs had the ability to take up Dil-Ac-LDL (Fig. 4A) and form capillary-like structures after 4 hours or 12 hours of incubation on Matrigel (Fig. 4B). To further demonstrate that PSCs-derived ECs have the ability to home to the vasculature of zebrafish in vivo (Orlova et al. 2014), we examined the engraftment potential of PSCs-derived ECs through the brief outline (Fig. 4C). When PSCs-derived ECs were injected into the vessels of 48 hpf zebrafish embryos, the cells were capable of integrating into the vascular system that had already developed (Fig. 4D). In addition, we performed cell therapy on zebrafish embryos with vasculature deficiency and less blood flowing through vessels after Sugen 5416 treatment. The results demonstrated that the percentage of zebrafish returning to normal was higher (26.71±5.86%) with PSCs-derived ECs treatment than with medium treatment (12.06±4.49%) (Fig. 4E). This finding suggested that the PSCs-derived ECs have potential for vascular repair.

**Fig. 4.**
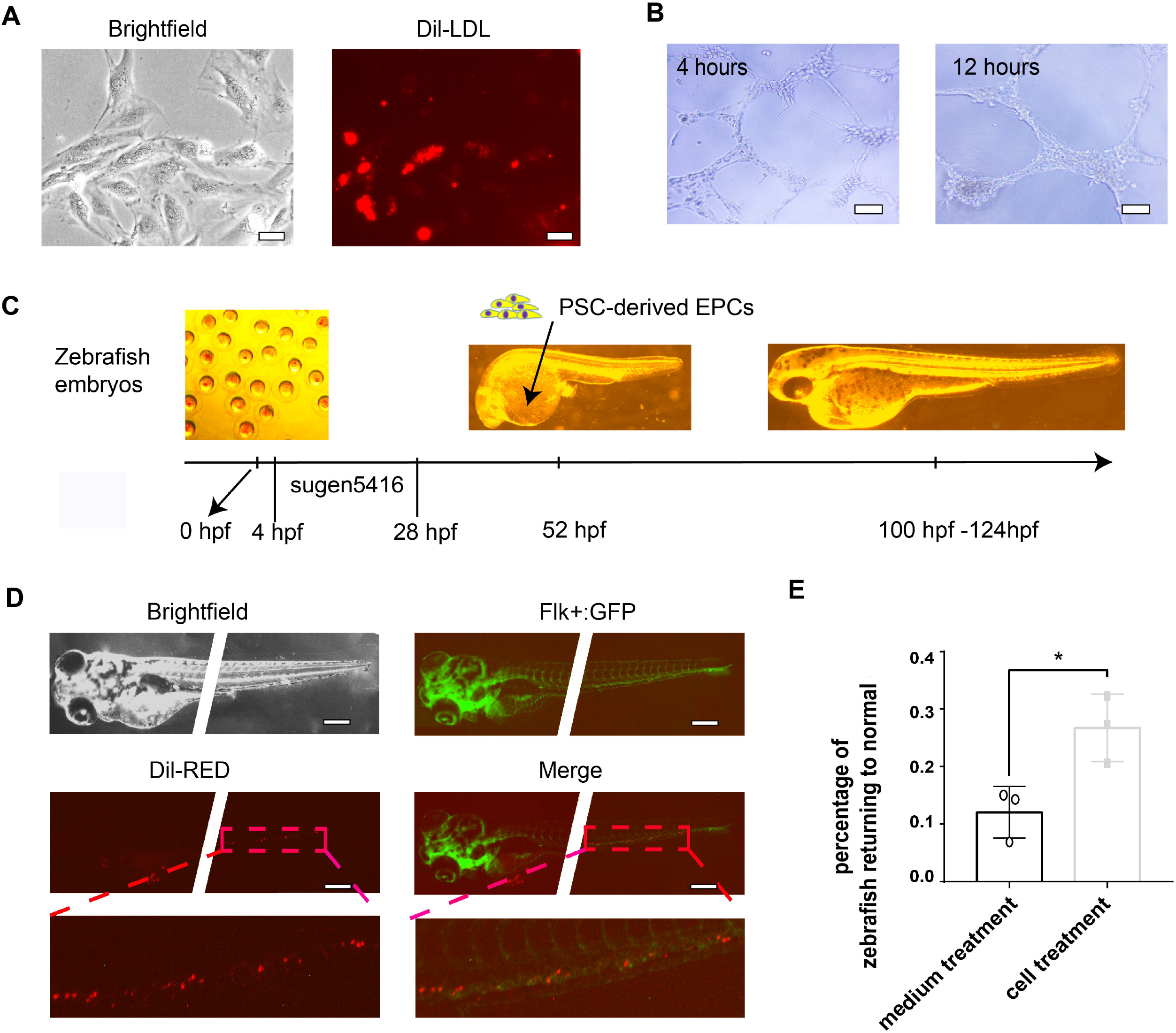
PSCs-derived ECs have functional characteristics of endothelial cells. (A) Uptake assay of Dil-acetylated LDL (scale bars: 50 μm). (B) Tube formation assay of H1-derived ECs after 4 hours and 12 hours (scale bars: 500 μm). (C) Brief outline of Zebrafish experiments. (D) Vascular competence of PSCs-derived ECs in a zebrafish xenograft model. Representative image of ECs-derived vessel-type structures (in red) within embryonic zebrafish (Flk: GFP; in green) 2 days after implantation, with magnification of the vessel. Scale bars are 300 μm. (E) Using a zebrafish model for gene therapy. Data are represented as the mean ± SD, n=3, and the experiments were repeated three times with 20-30 fish per condition. Student’s t test was performed (*p < 0.05).

### 3.5 PSCs-derived ECs have characteristics of EPCs

EPCs have two distinct EPC subtypes, early EPCs and late EPCs (also called endothelial colony-forming cells, ECFCs), which can be generated by the induction of ESCs/iPSCs; furthermore, high proliferation capacity of differentiated CD31^+^NRP1^+^ ECFCs is positively correlated with the expression level of NRP1 (Prasain et al. 2014; Yoder et al. 2007). To determine whether PSCs-derived ECs bear the property of ECFCs, we compared the cell morphology of PSCs-derived ECs with that of peripheral blood-derived EPCs and found that they were both typical cobblestone EC-like cells (Fig. 5A). We further detected increased expression of the NRP1 gene in PSCs-derived ECs during differentiation (Fig. 5B), which is consistent with ECFCs (Prasain et al. 2014). In addition, we also detected higher expression of another early EPC marker gene, *CD133* (*PROM1*) (Kanayasu-Toyoda et al. 2016), on day 7 by qRT-PCR (Fig. 5C). We also examined the expression levels of another two marker genes, *EFNB2* for arterial endothelium and *EPHB4* for vein endothelium, on day 7 by qRT-PCR. We found that 202-iPSC-ECs isolated on day 7 had a higher expression level of *EFNB2* than *EPHB4* (Fig. 5C). From the results demonstrated above, we reasoned that the PSCs-derived ECs have characteristics of EPCs and thought them as PSCs-derived EPCs.

**Fig. 5.**
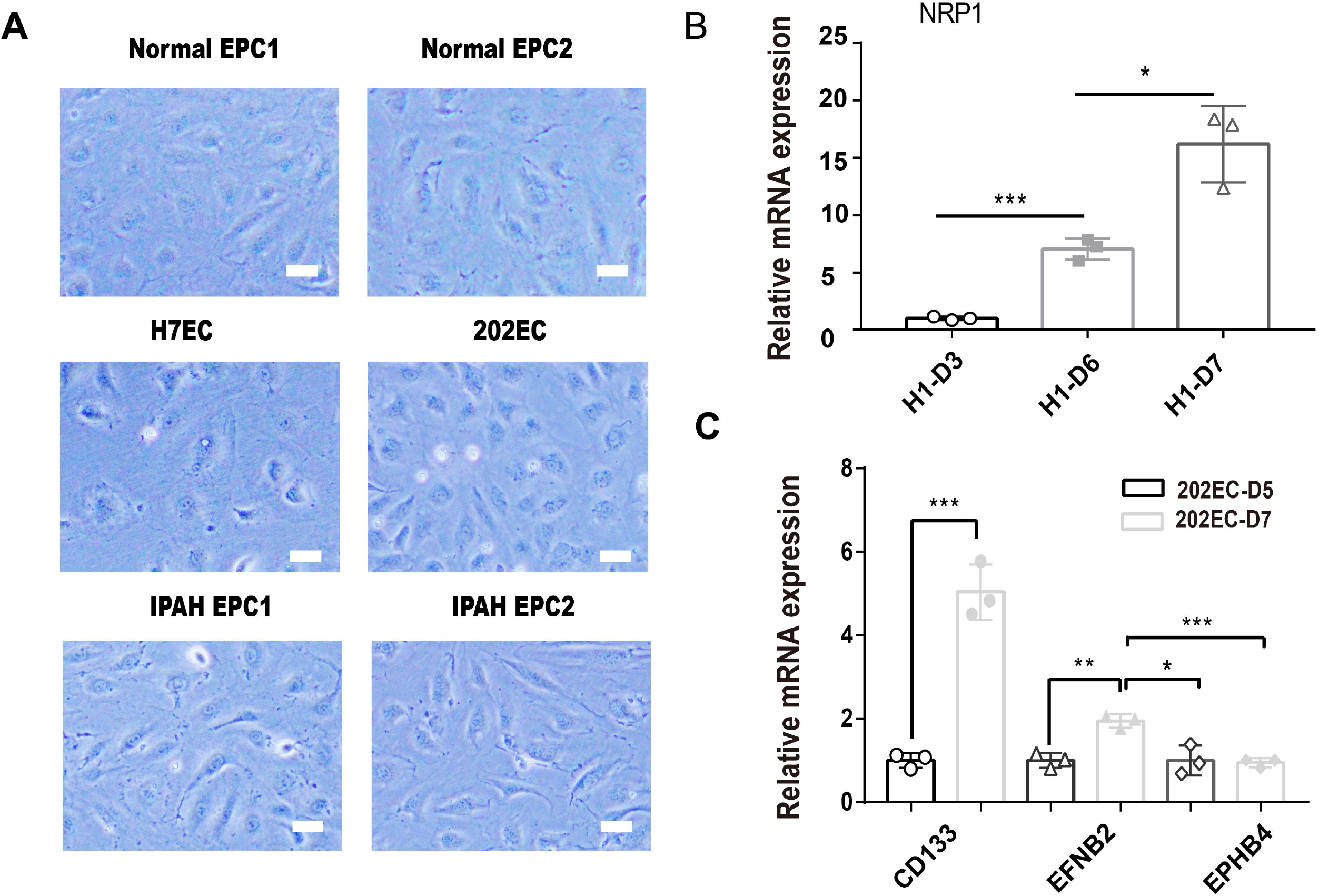
PSCs-derived ECs have characteristics of EPCs. (A) Comparison of the cell morphology of PSCs-derived ECs (H7EC, 202EC) with peripheral blood-derived normal EPCs (normal EPC1, normal EPC2) and IPAH-EPCs (IPAH EPC1, IPAH EPC2) (scale bars: 50 μm). (B) qRT-PCR of *NRP1*, which was reported to promote ECFC proliferation. *GAPDH* was used as an internal control. (C) ECs from DAY 5 or DAY 7 were analysed by the expression of the genes *CD133*, *EFNB2*, and *EPHB4. GAPDH* was used as an internal control. Data are represented as the mean ± SD, n=3, and the experiments were repeated three times. Student’s t test was performed (* p<0.05, **p < 0.01, ***p < 0.001).

### 3.6 Microarray and RNA-seq data further reveal the molecular characteristics and application potential of PSCs-derived EPCs

We next conducted bioinformatics analysis on datasets of PSCs-derived EPCs, normal EPCs, idiopathic pulmonary arterial hypertension (IPAH)-derived EPCs and GSE93511 (2D_MG_H1EC and 2D_MG_HUVEC) (Zhang, Schwartz, et al. 2017). First, we analysed in detail the expression patterns of key factors such as *CD31 (PECAM1), CD146 (MCAM), vWF, CD43(SPN), CD45(PTPRC), CD133(PROM1), KDR, NRP1, EFBN2, EPBH4* and *CD34*. The heatmaps showed that *PROM1, SPN and PTPRC* were not expressed in HUVECs, and had low expression in PSCs-derived EPCs, normal EPCs, IPAH-EPCs (Fig. 6A and 6B); moreover, *EFNB2* also had higher expression in PSCs-derived EPCs than normal EPCs and IPAH-EPCs (Fig. 6B), with the positive control showing higher level in 2D-MG-H1EC than 2D_MG_HUVEC (Fig. 6A). These results further confirmed that the PSCs-derived EPCs had features of arterial EPCs.

**Fig. 6.**
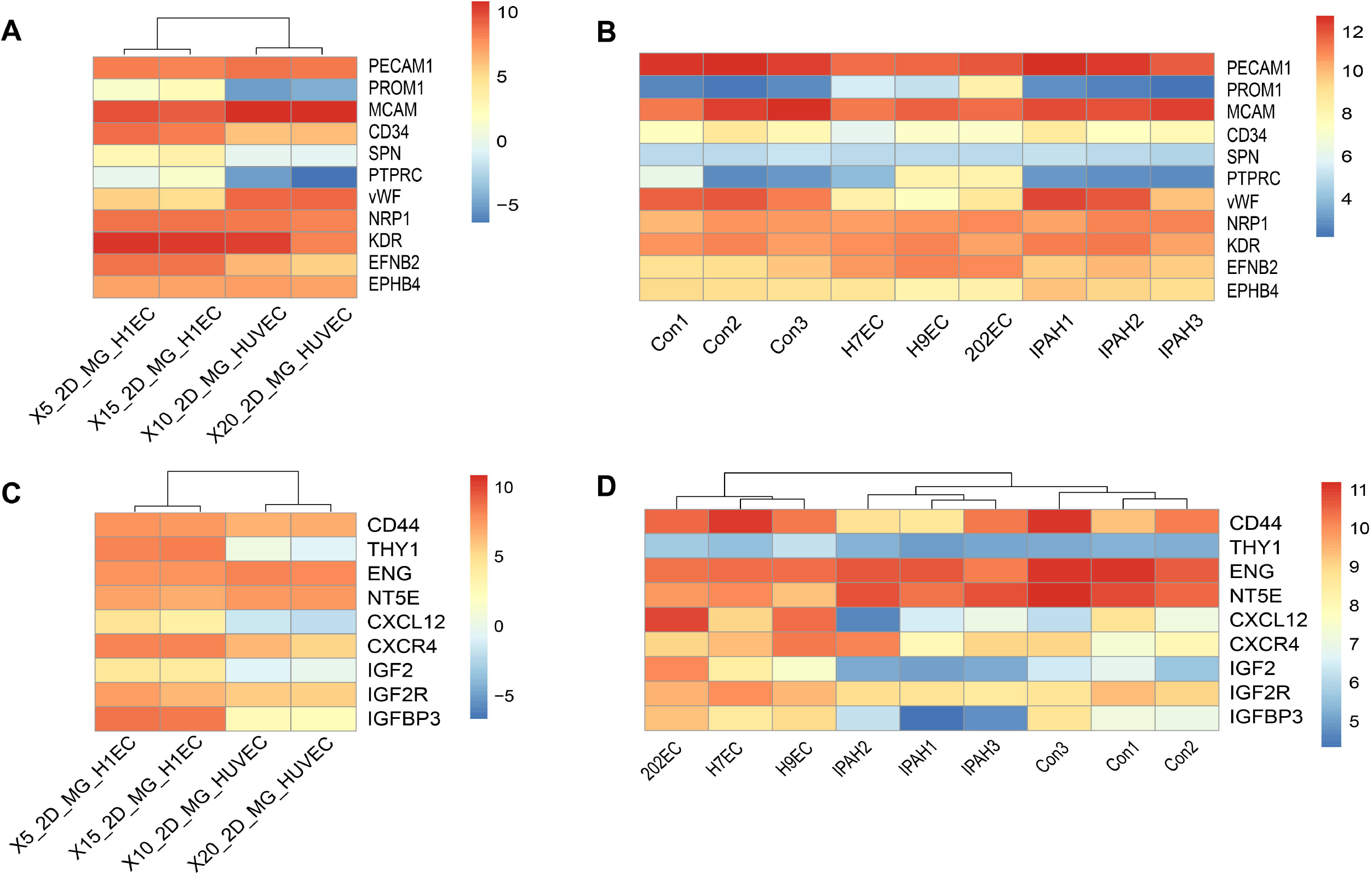
Bioinformatics analysis further reveals the characteristics of PSCs-derived EPCs. (A) and (B) Heatmap of EC-related genes from the 2D_MG_H1EC and 2D_MG_HUVEC datasets from GSE93511 and our microarray data in IPAH-EPCs (IPAH1, IPAH2, IPAH3), normal EPCs (Con1, Con2, Con3) and PSCs-derived EPCs (H7EC, H9EC, 202EC). *PROM1, SPN* and *PTPRC* were not expressed, and *EFNB2* had a higher expression level in HUVECs (A), but *PROM1, SPN* and *PTPRC* were highly expressed in PSCs-derived EPCs, normal EPCs, IPAH-EPCs and 2D-MG-H1EC (B). Moreover, *EFNB2* also had a higher expression level in PSCs-derived EPCs than in normal EPCs and IPAH-EPCs (B). (C) and (D) Heatmap of homing-related genes from the 2D_MG_H1EC and 2D_MG_HUVEC datasets from GSE93511 and our microarray data in IPAH-EPCs (IPAH1, IPAH2, IPAH3), normal EPCs (Con1, Con2, Con3) and PSCs-derived EPCs (H7EC, H9EC, 202EC). *IGF2, CXCL12* and *CD90(THY1)* were not expressed in HUVECs (C), but had higher expression levels in PSCs-derived EPCs, normal EPCs, IPAH-EPCs (D) and 2D-MG-H1EC (C). *IGFBP3* had a lower expression level in IPAH-EPCs than in PSCs-derived EPCs and normal EPCs (D).

Furthermore, we generated additional heatmaps of homing-related genes, such as *CXCR4* (Yuan et al. 2018), *IGF2/IGF2R*, and *CXCL12* (Zhuang et al. 2009; Xiaowei et al. 2013; Ferrari et al. 2011), and protein markers of mesenchymal stem cells, including CD73 (NT5E), CD44, CD90 (THY1), CD105 (ENG) (Caplan 2015; Jiang et al. 2019), and insulinlike growth factor-binding protein 3 (IGFBP3), which can improve vessel repair (Lofqvist et al. 2007; Chang et al. 2007). Heatmaps showed that *IGF2, CXCL12* and *CD90* (*THY1*) were not expressed in HUVECs (Fig. 6C) but had higher expression in PSCs-derived EPCs, normal EPCs, IPAH-EPCs and 2D-MG-H1EC (Fig. 6C-D). The rest of the genes had considerable expression levels in all cells, while we found that *IGFBP3* had lower expression in IPAH-EPCs than in PSCs-derived EPCs and normal EPCs (Fig. 6C-D). In addition, Gene Ontology (GO) analysis revealed that the genes highly expressed in PSCs-derived EPCs were related to the biological process of mesenchyme development, mesenchymal cell differentiation and mesenchyme morphogenesis (Fig. 7C). From the above, we think that PSCs-derived EPCs have the characteristics of mesenchymal stem cells for homing.

Next, we analysed the correlation between PSCs-derived EPCs and normal EPCs, and the results showed that the correlation was greater than 90% (Fig. 7A). In the process of cell culture, we found that IPAH-EPCs proliferated faster than PSCs-derived EPCs, and PSCs-derived EPCs grew slightly faster than normal EPCs (data not shown). From microarray data, we also found that the relative expression level of *MKi67*, a cell proliferation marker gene, was the highest in IPAH-EPCs, followed by PSCs-derived EPCs, and it was the lowest in normal EPCs (Fig. 7B). In view of the high similarity between PSCs-derived EPCs and normal EPCs, we performed Gene Ontology biological process (GOBP) analyses of PSCs-derived EPCs and IPAH-EPCs relative to normal EPCs. Our results showed that all genes upregulated in IPAH-EPCs were enriched in the top 20 biological processes related to cell division, while some genes of PSCs-derived EPCs were clustered in biological processes related to cell division, extracellular matrix, differentiation and development (Fig. 7C-D). These data again showed that IPAH-EPCs had a higher proliferation rate, which was consistent with previous results. In addition, to better understand the molecular characteristics of PSCs-derived EPCs, we preformed further GOBP analyses of PSCs-derived EPCs and normal EPCs relative to IPAH-EPCs. In the top 20 biological processes, our results showed that genes upregulated in PSCs-derived EPCs were enriched in immune-related biological processes, the top four of which are related to neutrophils (Fig. 7E); furthermore, some biological processes of normal EPCs were also related to immunity (Fig. 7F). This revealed that IPAH-EPCs had fewer immune-related molecular characteristics and enhanced proliferation capacity, suggesting that PSCs-derived EPCs maintained normal endothelial cell characteristics as well as function in therapy.

**Fig. 7.**
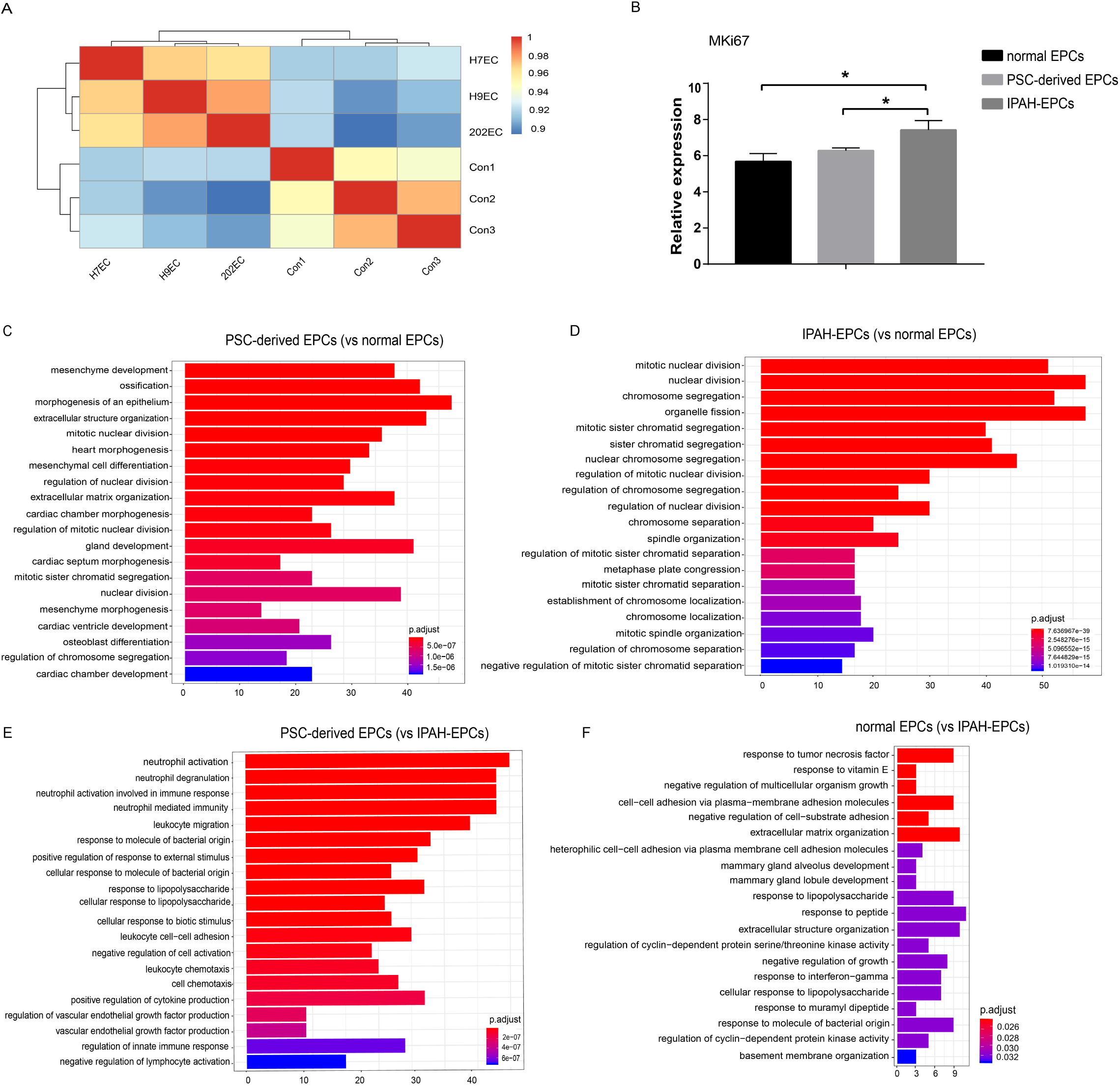
PSCs-derived EPCs have special molecular characteristics compared with normal EPCs and IPAH-EPCs (IPAH1, IPAH2 and IPAH3) based on our microarray data. (A) Correlation analysis between normal EPCs (Con1, Con2, Con3) and PSCs-derived EPCs (H7EC, H9EC, 202EC). (B) MKi67 relative expression from microarray data. Data are represented as the mean ± SD. n=3, Student’s t test was performed (*p < 0.05). (C) and (D) GOBP analysis (TOP20) of PSCs-derived EPCs and IPAH-EPCs relative to normal EPCs. (E) and (F) GOBP analysis (TOP20) of PSCs-derived EPCs and normal EPCs relative to IPAH-EPCs.

## 4. Discussion

Compared with other methods, such as EBs, coculture and monolayers, our optimized differentiation method is simplified and feasible for large-scale production. Although the differentiation efficiency of cells has been reported to be up to 48.43%-66.01% and 42.21%-51.9% by using coculture and EB methods respectively (Prasain et al. 2014), the process of differentiation is complex and time-consuming. The monolayer cell differentiation method is applied to derive endothelial cells because of its high efficiency and short cycle. For example, a small molecule differentiation approach shorten the process and induces more than 50% hPSCs into CD34^+^CD31^+^ cells in 19-9-11 iPSCs in 5 days (Lian et al. 2014); after 5 days of differentiation, H1 ESCs generate 74% ± 2% endothelial cells(Zhang, Schwartz, et al. 2017); then, the authors developed a “five factor” differentiation system (consisting of FGF2, VEGFA, SB431542, RESV, and L690) with which the maximum differentiation efficiency of endothelial cells reached 96.5% in 6 days (Zhang, Chu, et al. 2017). However, some specific conditions of endothelial differentiation still need to be optimized; for example, in a long-term process of endothelial cell differentiation, we could not achieve the same differentiation efficiency with a cell density of 1×10^5^ cells/cm^2^ (Zhang, Schwartz, et al. 2017). In addition, to obtain maximum differentiation efficiency, the conditions must be as consistent as possible in the differentiation system. The cell status, especially at the seeding stage, had a fundamental impact on the final efficiency. For example, using embryonic stem cells that have already differentiated greatly reduces the outcome production of ECs.

In addition, DMSO impacts PSCs-derived EPC differentiation during the whole process, which we previously ignored because it was only deemed as solvent to dissolve the chemicals. In our study, we used two chemical molecules, Y27632 and SB431542, although others have used CHIR99021 to promote EC differentiation (Nguyen et al. 2016; Zhang, Schwartz, et al. 2017). According to our experience, DMSO-treated cells need to be kept in a separate 37° C and 5% CO2 incubator, which could avoid inducing the differentiation of other cultured cells, suggesting that DMSO could affect cell differentiation and should be used with care. To better understand the function of DMSO, we searched papers referring to ESC differentiation, and found some reported that the efficiency of hPSC differentiation is improved even if hPSCs are treated with DMSO for a short time (Li et al. 2018); the addition of DMSO also downregulates the expression of stemness-related genes such as *OCT4* and *NANOG* and increases the proficiency of hepatic differentiation (Czysz, Minger, and Thomas 2015); DMSO at 0.01% and 0.1% concentrations can act as an induction agent for the formation of mesodermal phenotypes (Pal et al. 2012). These findings show that DMSO acts as a differentiation inducer in mesoderm differentiation.

The role of the inhibitor Y27632 in the process of PSCs-derived EC and HE differentiation has also been neglected because robust differentiation protocols started with single cells by using Y27632 at seeding to promote cell survival. This inhibitor can phosphorylate and activate the myosin II pathway and inhibit the E-cadherin-dependent apoptotic pathway (Vernardis et al. 2017). Some studies have reported that Y27632 enhances the differentiation of human term placenta-derived trophoblasts in vitro (Motomura et al. 2017), differentiation induction (Kurosawa 2012) and mesendodermal differentiation (Maldonado et al. 2016). In addition, the extension of treatment time to 3 days in the first stage of mesoderm induction may be able to substantially promote mesodermal cell production, consistent with another study (Nguyen et al. 2016).

PSCs-derived EPCs have special molecular characteristics and can be thought as a new EC-like subpopulation. Haematopoietic and endothelial progenitor cells share a number of cell-surface markers because they may originate from a common precursor, the haemangioblast (Ingram et al. 2004). It has been reported that CD34^+^ CD133^+^ VEGFR2^+^ cells are haematopoietic and may not actually be true EPCs (Medina et al. 2010), so we chose the endothelial marker CD31^+^ rather than CD34^+^ CD133^+^ VEGFR2^+^ to isolate PSCs-derived EPCs and then found their cell morphology as same as that of blood-derived normal EPCs; moreover, the expansion ability of PSCs-derived EPCs was similar with that of blood-derived normal EPCs, and they were both less proliferative than EPCs from IPAH patients. Finally, gene expression analysis from our microarray data showed that PSCs-derived EPCs expressing *CD40, CD90* and *CD105* were similar to mesenchymal stem cells, and the zebrafish transplantation experiment proved their function in vascular repairment; moreover, genes highly expressed in PSCs-derived EPCs were found to be involved in the biological processes of heart morphogenesis, cardiac chamber morphogenesis, cardiac septum morphogenesis, cardiac ventricle development and cardiac chamber development (Fig. 7C), which indicates that PSCs-derived EPCs have potential application value.

In the process of cell culture, we found that the cell morphology of IPAH-EPCs were not different from those of normal EPCs (Fig. 5A), but the proliferation ability of IPAH-EPCs was higher than that of normal EPCs, which is obvious in cell culture. Then, we analysed the expression levels of CD31, KDR, NRP1 and CD34 by flow cytometry and found that the percentage of NRP1^+^ cells was higher in IPAH-EPCs and PSCs-derived EPCs (Supplementary Fig. 2). Microarray analysis showed that the expression of the *IGFBP3* gene in IPAH-EPCs was significantly lower than that in normal EPCs and PSCs-derived EPCs (Fig. 6D). Moreover, GO clustering analysis revealed that IPAH-EPCs had high proliferative capacity and defects in immune-related gene expression (Fig. 7D and F), which could be further studied.

What is worth to mention is that we need focus on the most specific endothelial marker VE-cadherin (CDH5/CD144) used to isolate endothelial cells in other articles (Patsch et al. 2015; Gu et al. 2017). We also checked the expression level of *VE-cadherin*, and found that *VE-cadherin* has a higher expression level on day 5 than on day 7 in our optimized protocol by qRT-PCR (Supplementary Fig. 3A); in addition, another article reported that *CD31* and *VE-cadherin* have the similar expression tendency in the process of EC differentiation (Nguyen et al. 2016), so we chose to follow the original protocol with CD31 and CD34 (Zhang, Schwartz, et al. 2017). At last, we found that *VE-cadherin* expressed highly through microarray analysis (Supplementary Table 2), detected high expression of CD133 through immunofluorescence assay (Supplementary Fig. 3B) in isolated CD31^+^ cells which can form colonies (Supplementary Fig. 3C). These evidences confirmed that isolated CD31^+^ cells were PSCs-derived EPCs.

Moreover, the components in our EC differentiation system is compatible with other CDM medium. To verify the system and reduce the cost, we tested another CDM formulation and repeated the protocol with high differentiation efficiency as well. On the one hand, the function of each factor in the differentiation process was re-evaluated; on the other hand, this study lays a foundation for future application. PSCs-derived EPCs may be a new subpopulation have characteristics of early EPCs and mesenchymal stem cells for homing and possess greater cell therapeutic potential. Finally, we demonstrated that IPAH-EPCs had higher proliferation ability than normal EPCs.

## Authors’ contributions

K.Z.Q. and J. Y. participated in the research design, conducted the experiments, performed the data analysis and wrote the manuscript. All authors reviewed, revised the final version and approved manuscript submission.

## Ethics approval and consent to participate

All experiments were performed in accordance with the principles of the China Zebrafish Resource Center and approved by the Research Ethics Committee of Peking Union Medical College. All animal procedures were carried out in the Zebrafish Laboratory of State Key Laboratory of Medical Molecular Biology, Institute of Basic Medical Sciences, Chinese Academy of Medical Sciences and Peking Union Medical College. All experimental studies using human samples comply with the Declaration of Helsinki and were approved by the local ethics committee (Institute of Basic Medical Sciences, Chinese Academy of Medical Sciences and Peking Union Medical College). All persons gave informed consent before the study.

## Consent for publication

Not applicable.

## Acknowledgements

This research was supported by Grants from National Key Research and Development

Program of China-stem cell and translational research (No: 2016YFA0102300), CAMS Innovation Fund for Medical Sciences (CIFMS 2016-I2M-4-003), China National Thousand (Young) Talents Program of Jun Yang. The authors thank Hongtao Wang and Mengge Wang from State Key Laboratory of Experimental Hematology, Institute of Hematology & Blood Diseases Hospital, Tianjin 300020, China for their guidance with Hemogenic Endothelial (HE) differentiation.

